# Parachute geckos free fall into synonymy: *Gekko* phylogeny, and a new subgeneric classification, inferred from thousands of ultraconserved elements

**DOI:** 10.1101/717520

**Authors:** Perry L. Wood, Xianguang Guo, Scott L. Travers, Yong-Chao Su, Karen V. Olson, Aaron M. Bauer, L. Lee Grismer, Cameron D. Siler, Robert G. Moyle, Michael J. Andersen, Rafe M. Brown

**Affiliations:** Biodiversity Institute and Department of Ecology and Evolutionary Biology, University of Kansas, Lawrence, KS, 66045, USA; Department of Biological Sciences & Museum of Natural History, Auburn University, Auburn, Alabama 36849, USA; Chengdu Institute of Biology, Chinese Academy of Sciences, Chengdu, 610041, China; Department of Biomedical Science and Environmental Biology, Kaohsiung Medical University, Kaohsiung City, 80708, Taiwan; Department of Biology, 147 Mendel Hall, Villanova University, Villanova, PA 19085, USA; Herpetology Laboratory, Department of Biology, La Sierra University, Riverside, CA, 92515, USA; Department of Biology and Sam Noble Oklahoma Museum of Natural History, University of Oklahoma, Norman, OK 73072-7029, USA; Department of Biology and Museum of Southwestern Biology, University of New Mexico, Albuquerque, NM 87131, USA

**Keywords:** *Luperosaurus*, *Ptychozoon*, Phylogenomics, Species tree, Subgenera, Taxonomic vandalism, Ultraconserved elements

## Abstract

Recent phylogenetic studies of gekkonid lizards have revealed unexpected, widespread paraphyly and polyphyly among genera, unclear generic boundaries, and a tendency towards the nesting of taxa exhibiting specialized, apomorphic morphologies within geographically widespread “generalist” clades. This is especially true in the Australasia, where the monophyly of *Gekko* proper has been questioned with respect to phenotypically ornate flap-legged geckos of the genus *Luperosaurus*, the Philippine false geckos of the genus *Pseudogekko*, and even the elaborately “derived” parachute geckos of the genus *Ptychozoon*. Here we employ sequence capture targeting 5060 ultraconserved elements to infer phylogenomic relationships among 42 representative ingroup gekkonine lizard taxa. We analyzed multiple datasets of varying degrees of completeness (10, 50, 75, 95, and 100 percent complete with 4715, 4051, 3376, 2366, and 772 UCEs, respectively) using concatenated maximum likelihood and multispecies coalescent methods. Our sampling scheme was designed to address four persistent systematic questions in this group: (1) Are *Luperosaurus* and *Ptychozoon* monophyletic and are any of these named species truly nested within *Gekko*? (2) Are prior phylogenetic estimates of Sulawesi’s *L. iskandari* as sister to Melanesian *G. vittatus* supported by our genome-scale dataset? (3) Is the high elevation *L. gulat* of Palawan Island correctly placed within *Gekko*? (4) And, finally, where do the enigmatic taxa *P. rhacophorus* and *L. browni* fall in a higher-level gekkonid phylogeny? We resolve these issues; confirm with strong support some previously inferred findings (placement of *Ptychozoon* taxa within *Gekko;* the sister relationship between *L. iskandari* and *G. vittatus*); resolve the systematic position of unplaced taxa (*L. gulat*, and *L. browni*); and transfer *L. iskandari, L. gulat, L. browni*, and all members of the genus *Ptychozoon* to the genus *Gekko*. Our unexpected and novel systematic inference of the placement of *Ptychozoon rhacophorus* suggests that this species is not related to *Ptychozoon* or even *Luperosaurus* (as previously expected) but may, in fact, be most closely related to several Indochinese species of *Gekko*. With our final, well-supported topologies, we recognize seven newly defined subgenera to accommodate ∼60 species within the more broadly defined and maximally-inclusive Australasian genus *Gekko*. The newly defined subgenera will aide taxonomists and systematists in species descriptions by allowing them to only diagnose putatively new species from the most relevant members of the same subgenus, not necessarily the phenotypically variable genus *Gekko* as a whole, and we argue that it appropriately recognizes geographically circumscribed units (e.g., a new subgenus for a novel clade, entirely endemic to the Philippines) while simultaneously recognizing several of the most systematically controversial, phenotypically distinct, and phylogenetically unique lineages. An added benefit of recognizing the most inclusive definition of *Gekko*, containing multiple phylogenetically-defined subgenera, is that this practice has the potential to alleviate taxonomic vandalism, if widely adopted, by creating formally available, supraspecific taxa, accompanied by character-based diagnoses and properly assigned type species, such that future, more atomized classifications would necessarily be required to adopt today’s subgenera as tomorrow’s genera under the guidelines of The Code of Zoological Nomenclature. Not only does this simple practice effectively eliminate the nefarious motivation behind taxonomic vandalism, but it also ensures that supraspecific names are created only when accompanied by data, that they are coined with reference to a phylogenetic estimate, and that they explicitly involve appropriate specifiers in the form of type species and, ultimately, type specimens.

## 1. Introduction

The family Gekkonidae is the largest gekkotan lizard family comprising ∼1200 species and 61 genera (Uetz et al., 2018). Within this family, the genus *Gekko* (Uetz et al., 2018) contains 59 currently recognized species, and the allied but phenotypically distinct species of Flap-legged (*Luperosaurus*) and Parachute (*Ptychozoon*) geckos, each with 13 recognized species have been the subject of several recent phylogenetic analyses (Brown et al., 2012a, b; Grismer et al., 2018, in press). The results of these studies suggest that the genus *Gekko* is rendered paraphyletic by some species of the other two, highly autapomorphic genera (Brown et al., 2012a, b; Heinicke et al., 2012; Pyron et al., 2013). The enigmatic phylogenetic relationships of these peculiar geckos have been the focus of traditional character-based classifications (Boulenger, 1885; Wermuth, 1965), analyses of external and internal anatomical morphological characters (Brown et al., 2001), and more recent multi-locus Sanger sequence datasets with nearly complete taxon sampling, save a few key taxa, such as the secretive *Luperosaurus browni* (Brown et al., 2012a, b). To date, analyses of Sanger datasets have inferred fairly consistent topologies for the genus *Gekko*; however, phylogenetic uncertainty increased with the inclusion of closely related genera. For example, lineages of *Luperosaurus* not only appear to render *Gekko*, but also *Lepidodactylus* (not considered here), paraphyletic (Heinicke et al., 2012; Oliver et al., 2018). To date, phylogenetic analyses that include taxa outside of *Gekko* have led to unclear taxonomic boundaries, nomenclatural instability, and frequent transfers of taxa between genera (Russell, 1979; Rösler et al., 2012; Brown et al., 2000, 2007, 2012a, b). Thus, a well-resolved phylogeny of *Gekko* and its allies remains outstanding.

To address higher-level relationships of *Gekko sensu stricto* across its wide distribution, we apply a genomic approach targeting 5060 ultraconserved elements (UCEs) and modern phylogenomic analyses to address the following questions: (1) Are *Luperosaurus* and *Ptychozoon* each monophyletic, and are they or any of their included taxa nested within *Gekko*? (2) Does our surprising former estimate of the systematic position of Sulawesi’s *L. iskandari* (sister to Melanesian *G. vittatus*), hold up under phylogenomic inference? (3) Is the high elevation Palawan Island species “*Luperosaurus*” *gulat* correctly placed in *Gekko* (represented by a single specimen and degraded accompanying DNA sample) as suggested by a previous analysis of two Sanger loci? (4) Finally, where do the elusive *P. rhacophorus* and *L. browni* fall within the higher-level gekkonid phylogeny?

## 2. Materials and methods

### 2.1. Taxon sampling

Ingroup sampling included 42 individuals representing two species of *Lepidodactylus*, two species of *Pseudogekko*, five *Ptychozoon* species, seven *Luperosaurus* species, and 25 species of *Gekko*, including *G. smithii*, which is the sister species to the generotype, *G. gecko* Linnaeus, 1758 (Table S1). *Cyrtodactylus baluensis, C. jellesme*, and *C. redimiculus*, were chosen as distant outgroup taxa to root the tree based on a recent phylogenetic study of Southeast Asia geckos (Brown et al., 2012b). Our taxon sampling was selected to resolve a polytomy consisting of several clades assigned to the genera *Gekko, Luperosaurus*, and *Ptychozoon* (Brown et al., 2012a). Issues relating to the relationships of some *Luperosaurus, Lepidodactylus*, and *Pseudogekko* taxa (Heinicke et al., 2012; Oliver et al., 2018) are the focus of a different study and will be discussed elsewhere.

### 2.2. Data collection and sequence capture of UCEs

Data collection and sequence capture of UCEs took place in two different batches. The first batch of samples was prepared under the following conditions: high quality genomic DNA was extracted from 26 individuals using the Qiagen DNAeasy^®^ kit following the animal tissue protocol. Genomic DNA concentrations were measured using a QUBIT^®^ 2.0 fluorometer and were standardized to 500 ng in 50 μL. We sheared genomic DNA using a Covaris S220 with the following setting: peak power 175 W, duty factor: 2.0%, cycles per burst: 200, duration: 45 seconds. We prepared Illumina libraries using a NEB/KAPA library preparation kits following Faircloth et al. (2012)–described in detail at http://ultraconserved.org. Next, we ligated universal iTru stubs (Glenn et al., 2016) in place of standard-specifics to allow for dual indexing. This was proceeded by a second 1X volume AMPure XP bead clean up after stub ligation. Followed by a 17-cycle PCR with NEB Phusion High-Fidelity PCR Master Mix of iTru Dual-indexes (Glenn et al., 2016) with the library fragments.

We then quantified the dual-indexes PCR product and library fragments using a Qubit 2.0 Fluorometer, pooled libraries into groups of eight, and enriched each pool for 5060 UCEs. We targeted UCEs with 5472 probes from the Tetrapod 5Kv1 probeset (Arbor Biosciences, formerly MYcroarray). See Faircloth et al. (2012) and http://ultraconserved.org/#protocols for details on probe design. Next, we amplified enriched pools using limited-cycle PCR (17 cycles) and sequenced our enriched libraries on a single lane of an Illumina HiSeq 2500 (PE100 reads) at the KU Genome Sequencing Core. The second batch of genomic extractions, for 20 additional individuals, used the Maxwell©RSC Tissue DNA kit on the Promega Maxwell©RSC extraction robot. Genomic DNA was quantified on a Promega Quantus™fluorometer and standardized to 1000 ng in 50 μL of ultrapure DNA-grade water. Quantified samples were outsourced to MYcroarray (now Arbor Biosciences) for library preparation, following the same sequence-capture protocol outlined above. Libraries were sequenced on a single lane of an Illumina HiSeq 3000 (PE150) at Oklahoma Medical Research Foundation (OMRF).

### 2.3. Data processing

Data were demultiplexed at the KU sequencing facility for the first batch of samples and at the OMRF for the latter batch. We subjected all samples to a custom bioinformatics pipeline version 1.0 (https://github.com/chutter/) to filter and remove adaptor contamination, assemble, and export alignments. We filtered samples and removed adapters by using the bbduk.sh script (part of BBMap/BBTools; http://sourceforge.net/projects/bbmap/) with the following parameters: ftm = 5, ktrim = r, k = 23, mink = 8, hdist = 1, tbo, tpe, and minlength = 25. Next we used the bbsplit.sh script to remove other sources of non-focal organism contamination (e.g., *Achromobacter*, *Acidaminococcus*, *Acinetobacter*, *Afipia*, *Agrobacterium*, *Alcaligenes*, *Aminobacter*, *Aspergillus*, *Bradyrhizobium*, *Brevundimonas*, *Burkholderia*, *Caenorhabditis*, *C. elegans, Corynebacterium, Curvibacter, E. coli, Flavobacterium, Haemophilus, Helcococcus, Herbaspirillum*, *Legionella*, *Leifsonia*, *Magnaporthe*, *Malassezia*, *Mesorhizobium*, *Methylobacterium*, *Microbacterium*, *Moraxella*, *Mycoplasma*, *Novosphingobium*, *Ochrobactrum, Pedobacter, Penicillium, Phyllobacterium, Pseudomonas, Pseudonocardia, Puccinia, Ralstonia, Rhodococcus, Saccharomyces, Salmonella, Scerevisiae, Schizosaccharomyces, Sphingomonas, Staphylococcus, Stenotrophomonas*, and the UniVec database for vector contamination) with a minid = 0.95, followed by additional decontamination of singleton reads. Following decontamination, we removed adaptors and error corrected using AfterQC (Chen et al., 2017) with parameters set to: qualified quality phred=0, number of base limit=10, sequence length requirement=35, unqualified base limit=60, trim front and tail bases set to zero, and applied to both reads. We merged paired end reads using bbmerge-auto.sh (part of BBMap/BBTools; http://sourceforge.net/projects/bbmap/) with the following settings: verystrict to decrease the merging rate, kmer length=60, extend reads by 60 if a failed merge attempt, with error correction (verystrict = t rem k = 60 extend2 = 60 ecct).

### 2.4. UCEs assembly, probe matching, alignment, and trimming

*De novo* assembly was conducted with SPAdes v3.11.1 (Bankevich et al., 2012) using multiple k-mer sizes (21, 33, 55, 77, 99, and 127), with the setting to expect significant amounts of gaps, and with haplotype assembly phasing. We used an array of k-mer values to aide in merging orthologous contigs resulting from different k-mer sizes. We further assembled contigs using the DIPSPADES (Safonova et al., 2015) function to better assemble exons and orthologous regions by generating a consensus sequence from both orthologous and haplotype regions. The dedupe.sh script (part of BBMap/BBTools; http://sourceforge.net/projects/bbmap/) was used to remove near exact duplicates and pblat (Kent, 2002; Meng, 2018) was used to match samples to targeted reference loci with a tile size set to eight and minimum sequence identity set to 60. All matching loci per species were then merged into a single file for downstream UCE alignments. Prior to UCE alignment, all loci recovered in only one sample were discarded. UCE alignments were constructed using the high accuracy option in MAFFT v7.130b (Katoh and Standley, 2013) with a max iteration of 1000, automatically adjust the reading direction, gap opening penalty set to three, and an offset “gap extension penalty” set to 0.123. Because large UCE alignments usually contain long stretches of poorly aligned sequence regions we internally trimmed all of the alignments using trimaAL (Capella-Gutiérrez et al., 2009). We applied the automated1 command which implements a heuristic search to choose the most appropriate mode based on the given characters. Following alignment trimming we generated five datasets with varying levels of loci completeness (100p, 95p, 75p, 50p, and 10p), where 95p requires 95 percent of the individuals in the alignment for a locus to be included. Summary data for all datasets were produced using scripts available at https://github.com/dportik/Alignment Assessment. Frequency distributions of the genomic data are presented on a per locus basis for 45 individuals (Fig. 1A-F) and all data files are deposited on DRYAD (XXXXXXXXXXXX) and raw sequence data is deposited on GenBank Sequence Read Archive (XXXXXXXXXX).

**Fig. 1.**
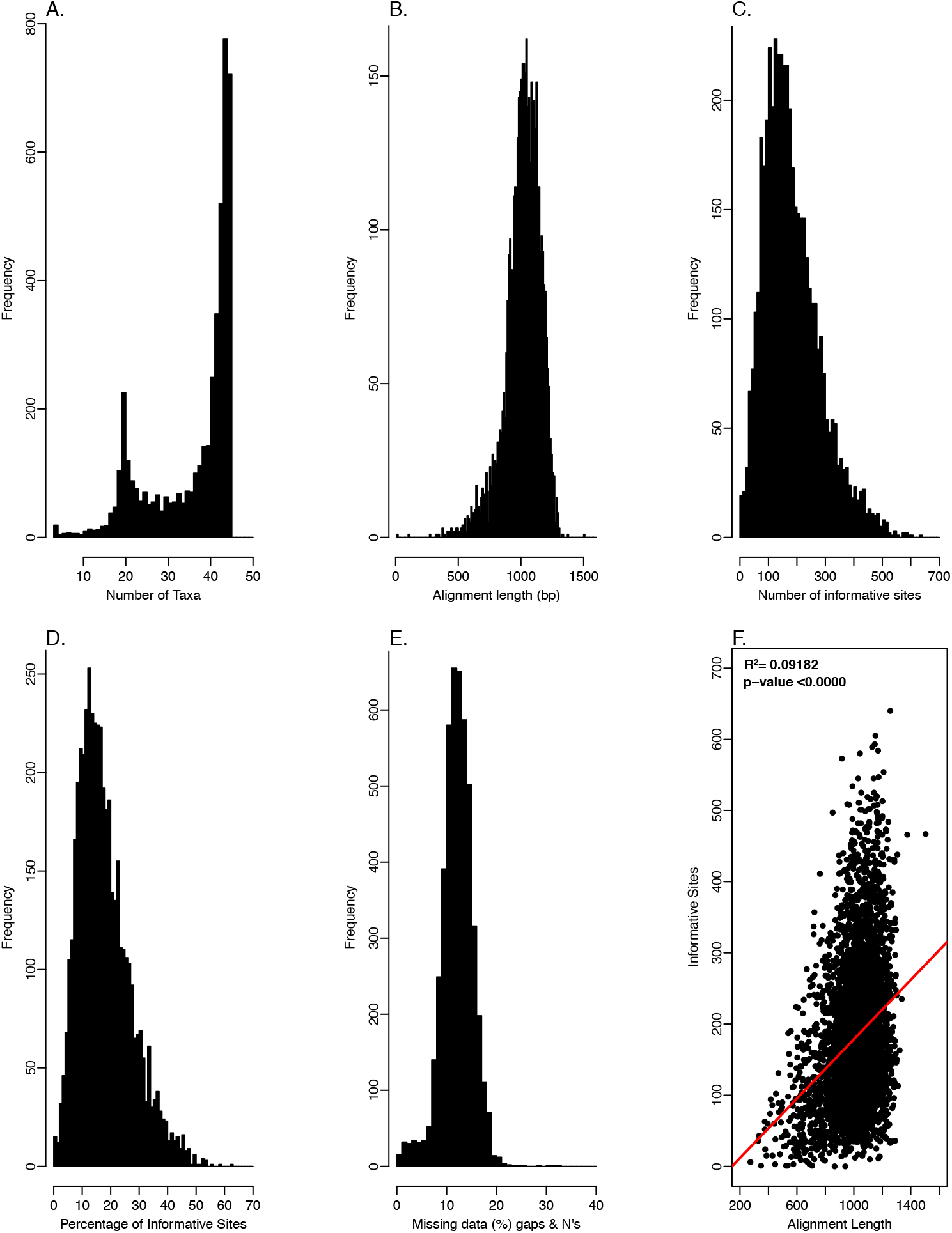
Frequency distributions of assembled loci from 4718 targeted loci. Summary statistics from the raw Illumina reads. A. Number of taxa present in the alignment. B. Alignment length. C. Number of informative sites. D. Percentage of informative sites. E. Percentage of missing data, gaps, and N’s present in the alignments. F. Alignment length with respect to informative sites; note, there is a significant positive correlation between alignment length and number of informative sites, R= 0.09182, p-value ≤ 0.0000.

### 2.5. Phylogenomic analyses

To reconstruct higher-level phylogenetic relationships of *Gekko*, we analyzed all five datasets (100p–10p) using maximum likelihood for the concatenated datasets, as well as species-tree analyses. We prepared concatenated PHYLIP files for each dataset with a single partition and estimated maximum likelihood (ML) phylogenies using IQ-TREE v1.6.7 (Nguyen et al., 2015), applying the GTR+**Γ** model of molecular evolution. We assessed nodal support using 1000 bootstrap pseudoreplicates via the ultrafast-bootstrap approximation algorithm (Minh et al., 2013). Nodes with Bootstrap (BS) ≥95 were considered to be well-supported (Minh et al., 2013; Wilcox et al., 2002).

Under certain conditions, gene tree-species tree methods have advantages over the analysis of concatenated datasets (Kubatko and Degnan, 2007; Edwards et al., 2007, 2016), but they may also be sensitive to missing data (Bayzid and Warnow, 2012) and to the resolution of individual gene trees (Castillo-Raímrez et al., 2010). Here we applied two different species tree methods: ASTRAL-III (Zhang et al., 2017) and SVDQuartets (Chifman and Kubatko 2014). For ASTRAL, we summarized individual gene trees as input files, whereas we provided a partitioned nexus file for SVDQuartets without estimating individual gene trees, *a priori*. Under both methods we analyzed 100p and 50p complete datasets. Individual gene trees for each locus were estimated in, IQ-TREE with implementing ModelFinder (Kalyaanamoorthy et al., 2017) to rapidly estimate an accurate model of evolution, which were used as the input trees for downstream species tree analyses in ASTRAL-III. We estimated quartet support values implemented in ASTRAL-III, where the quartet support values are the posterior estimate at a given branch where each gene tree quartet agrees with the respective branch.

Species tree analyses were inferred using SVDQuartets and implemented in an alpha-test version of PAUP* v4.0a15080 (Swofford, 2003). This algorithm randomly samples quartets using a coalescent model and a quartet amalgamation heuristic to generate a species tree and has proven useful and accurate for estimating species trees from complete alignments of large genomic datasets (Chou et al., 2015). All possible quartet scores were evaluated from the entire alignment using a multispecies coalescent tree model with 100 bootstrap replicates performed to calculate nodal support. The program Quartet MaxCut v2.1.0 (Snir and Rao, 2012) was used to construct a species tree from the sampled quartets.

## 3. Results

### 3.1. UCE sequencing, assembly and alignment

Following enrichment and sequencing, we obtained an average of 5462 contigs per sample (range = 2812–8756). An average (per sample) of 4718 of these contigs matched the UCE loci from capture probes (range = 2976–5277). The average length of UCE-matching contigs was 946 bp (range = 550–1413). For the dataset specifying no missing data, we recovered 772 UCE loci across 45 taxa (797,846 bp including indels) with 132,677 informative sites. For the 50p dataset, we recovered 4,051 UCE loci across 45 species, with average length of 4,171,817 bp (including indels). For the datasets allowing missing loci for any taxon, we recovered 4715 loci (4,794,263 bp) for 10p, 3376 loci (3,508,564 bp) for 75p, and 2366 loci (2,543,167 bp) for 95p UCE loci across 45 species. We provide summary statistics for sequencing and alignments in Table S1 and in Figure 1.

### 3.2. Phylogenomic analyses

For the 100p dataset of 772 UCE loci, the maximum likelihood analysis produced an optimal topology in which all but three nodes received over 95% bootstrap support (Fig. 2A). The ML topologies from the four other variably incomplete data matrices (4715, 4051, 3376, and 2366 loci, respectively) reflect the same relationships as the 100p dataset, with only a few nodes lacking strong support (Fig. 2A–E).

**Fig. 2.**
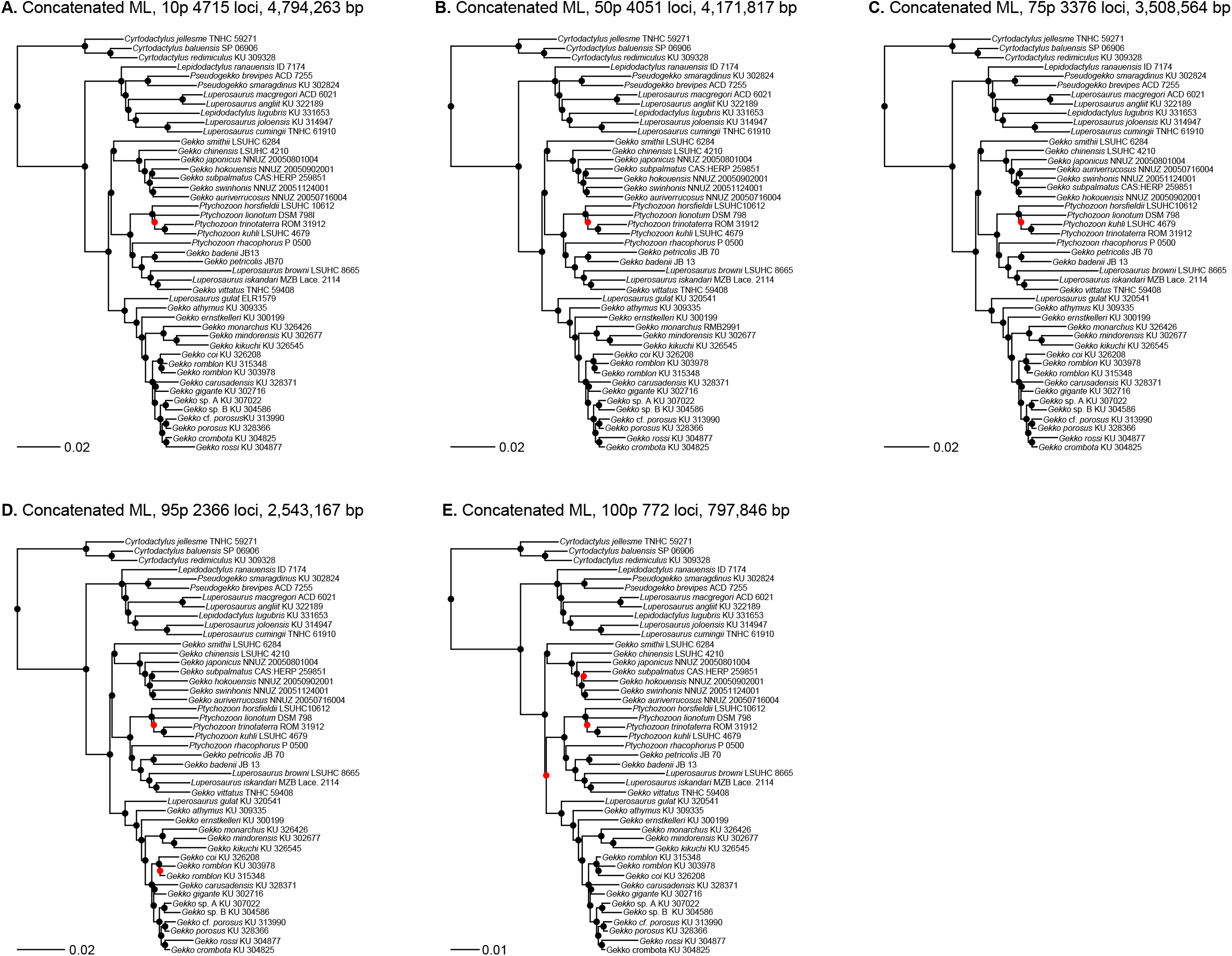
Maximum likelihood phylogenies inferred from concatenated datasets from IQ-TREE analyses. A. 10p 4,715 loci, 4,794,263 bp. B. 50p 4,051 loci, 4,171,817 bp. C. 75p 3,376 loci, 3,508,564 bp. D. 95p 2366 loci, 2,543,167 bp. E. 100p 772 loci, 797,846 bp. Black dots represent bootstrap support values greater than 95, and red dots are support values 94 and below.

Our phylogenomic estimate provided strong support for novel relationships, placing unequivocally several rare taxa of unknown affinities, while simultaneously resolving several unresolved, problematic relationships. Our UCE data resolved the three-clade polytomy of Brown et al. (2012a: Fig. 1, node 3)—all of the deeper internodes of *Luperosaurus* + *Gekko* + *Ptychozoon*—and most nodes in our concatenated trees were also resolved with strong support in our species tree analyses (Fig. 3).

**Fig. 3.**
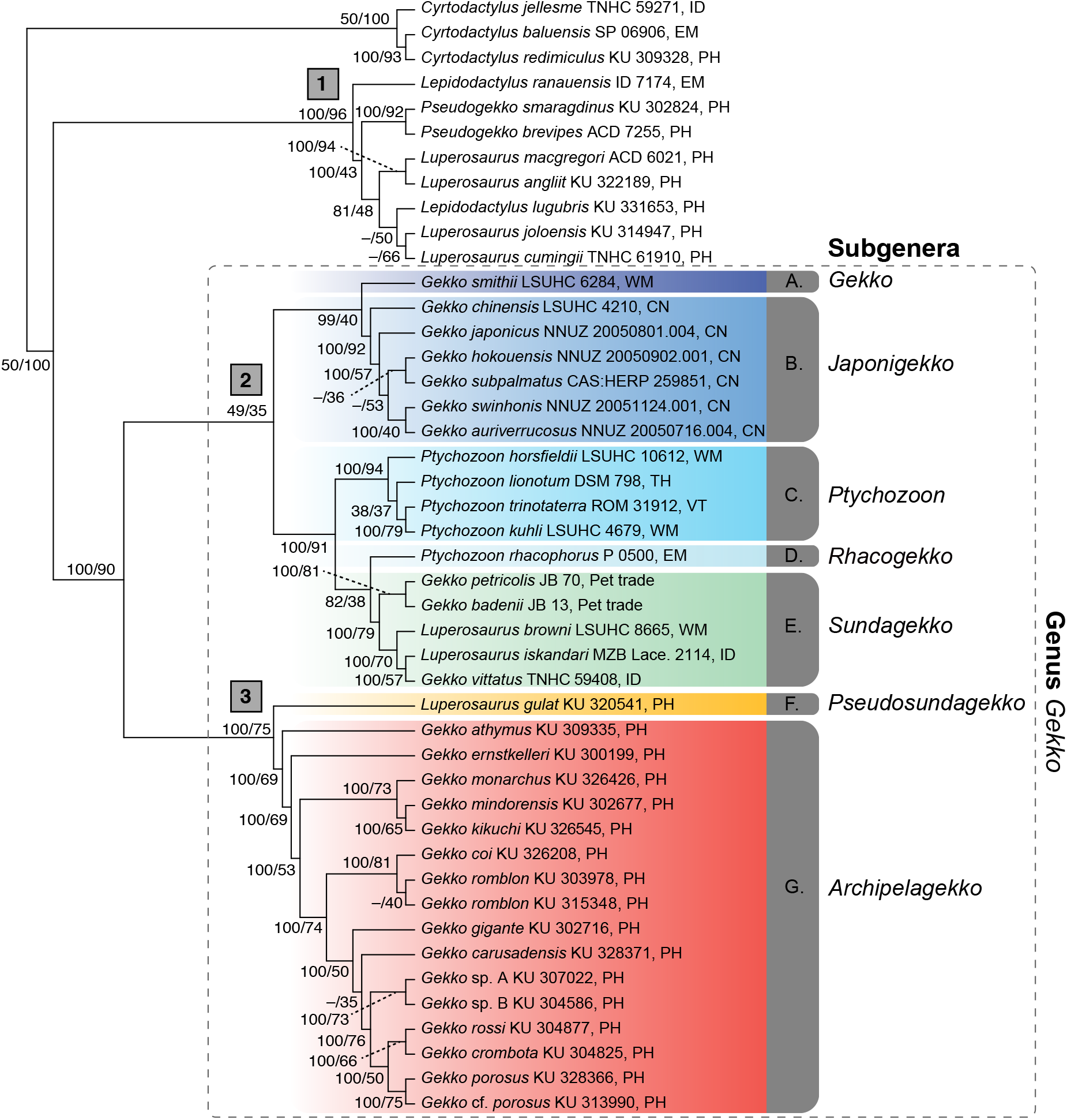
ASTRAL species tree of gekkonid species included in this study based on the alignment with no missing data 100p (722 loci, 797,846 bp). Node values are SVDQuartet support values and quartet support values inferred from ASTRAL-III, respectively. Bold numbers in grey boxes refer to the three large clades recovered from the phylogenetic analyses. All subgenera groups are labeled A–G and are colored according to the new subgeneric designations. Country codes are as follows: CN, China; EM, East Malaysia; ID, Indonesia; WM, West Malaysia; PH, Philippines; TH, Thailand; VT, Vietnam.

We inferred strong support for three large clades (Fig. 3, clades 1–3), the first of which contains taxa previously referred to *Luperosaurus* (including the generotype, *L. cumingii*, eliminating any doubt that this clade should be considered true *Luperosaurus*), *Lepidodactylus*, and *Pseudogekko*. Surprisingly, a close relationship between *Lepidodactylus lugubris* and members of true *Luperosaurus* (exemplified by *L. cumingii*) was well-supported in the concatenated analyses; however, this group received variable support in the species tree analyses (Fig. 3 vs. Fig. 2A–E; see also Heinicke et al., 2012). A second well-supported clade, including *Gekko smithii* (uncontroversial, previously documented sister to the genus *Gekko*’s generotype species, *G. gecko*; Rösler et al., 2011), comprises numerous species previously referred to *Gekko, Ptychozoon* (including its generotype species *P. kuhli*), and a subset of insular *Luperosaurus* taxa (Fig. 3).

The true *Gekko* (*+ Ptychozoon*) clade inferred in our phylogenomic analyses was recovered with strong support as most closely-related to the Philippine *Gekko* clade (Fig. 3, clade G), with one surprising result. In contrast to the two-locus analysis of Brown et al. (2012a), in which the newly-discovered, enigmatic high-elevation Palawan Island endemic *Luperosaurus gulat* was recovered as part of the *Gekko + Ptychozoon* clade, but with no support for a close relationship to any of its included taxa, *L. gulat* was recovered here as the first-diverging lineage among the nearly pure Philippine *Gekko* radiation (Figs. 2A–E, 3).

Similar to previous studies (small multi-locus datasets; Brown et al., 2012a; Heinicke et al., 2012), *Gekko* species are arranged in three major subclades: two *Gekko* clades and one *Gekko* + *Ptychozoon* + *Luperosaurus. Gekko* as a whole is non-monophyletic because of the inclusion of *Ptychozoon* and some *Luperosaurus* (Figs. 2A–E, 3), and remaining “*Luperosaurus*” taxa (species not recovered in the generotype group) are placed with strong support in four different clades (Figs. 2A–E, 3). The close relationship between the Southeast Asian species pair, *L. browni + L. iskandari*, and *Gekko vittatus* is strongly supported, and these are in turn related to *G. badenii* and *G. petricolis* (Fig. 3).

Finally, another novel result is the strongly-supported placement of *Ptychozoon rhacophorus* apart from the remaining species of *Ptychozoon*, but as sister to the (*G. badenii* + *G. petricolis*) ((*L. iskandari* + *Gekko vittatus*), *L. browni*) clade of Indochinese, Southeast Asian, and Southwest Pacific island taxa, respectively.

## 4. Discussion

### 4.1. Phylogeny, geographic regionalism, and biogeography

Our concatenated ML analyses of 772 (100p), 3,376 (75p), and 4,715 (10p) loci produced identical relationships and these results are similar, in some ways, to those of earlier analyses of few loci (Brown et al., 2012a; Heinicke et al., 2012). Nevertheless, despite the similarity of these results to the partially resolved earlier study of Brown et al. (2012a), our phylogenomic analysis of UCE-probed loci using both concatenated and species tree analyses produced novel relationships that allow us to make several strong conclusions. This study establishes the strongly supported placement of the formerly problematic taxon *Luperosaurus gulat* (Brown et al., 2010), provides resolution of a polytomy from an earlier study (Brown et al., 2012a), and confirms the nested placement of taxa with phenotypically apomorphic morphologies within a large clade characterized by a generalized suite of pleisiomorphic traits (Brown et al., 2012b; Heinicke et al., 2012). Aside from confirming the biogeographic regionalism of earlier inferred *Gekko* clades— recognized in our revised classification proposed here as subgenera (see below)—our phylogenomic results have additional biogeographic significance. With *L. gulat* (Brown et al., 2010) now recognized as a *Gekko* species that is strongly supported as sister to the remaining Philippine *Gekko*, ancestral range reconstruction of the Palawan Microcontinent Block at the base of the Philippine clade is considerably strengthened (Siler et al., 2012). The first three successively branching Philippine lineages (*L. gulat, G, athymus*, and *G. ernstkelleri*) are all Palawan Microcontinent Block endemics, as are several species nested more distally in this clade (*G. palawanensis, G. romblon, G. coi*, and, in part, *G. monarchus*). The phylogenetic placement of *Gekko gulat*, n. comb., thus adds to a suite of recent studies emphasizing the Palawan Microcontinent Block-origins of Philippine archipelago-wide endemic clades, ancient isolation of Eurasian lineages (to the exclusion of Sundaic lineages), a paleotransport-and-dispersal facilitated colonization scenario, and the pivotal role of the Palawan Ark mechanism for biogeographic contributions to accumulation of the archipelago’s land vertebrate faunal megadiversity (Blackburn et al., 2010; Siler et al., 2012; Brown et al., 2013, 2016; Grismer et al., 2016; Chan and Brown, 2017).

### 4.2. Evolution of phenotypic novelty

Our phylogenomic estimate of selected *Gekko* taxa both confirms previous findings of paraphyly with respect to *Ptychozoon* and some species of *Luperosaurus* (Brown et al., 2012a, b; Heinicke et al., 2012) and provides new insight into the evolution of morphological novelty within gekkonid lizards (Gamble et al., 2012; Oliver et al., 2018). Although *Gekko* is known to be a phenotypically variable, highly diverse clade (Rösler et al., 2011), with some clades more variable than others, none exhibit more structural body plan novelty than members of the genus *Ptychozoon* (Brown et al., 1997; Russell et al., 2001). Only *Luperosaurus iskandari* (Brown et al., 2000) approaches the degree of elaboration of dermal structures bordering the limbs, tail, and body—clear adaptations for directed aerial descent (parachuting, gliding; Heyer and Pongsapipatana, 1970; Marcellini and Keefer, 1976; Russell, 1979; Brown et al., 2001; Young et al., 2002; Heinicke et al., 2012) and camouflage (Barbour, 1912; Tho, 1974; Brown et al., 1997; Vetter and Boride, 1977). Within the family, evolution of similar structures capable of lift generation during aerial locomotion and breaking up the body’s outline when at rest on vertical surfaces have evolved multiple times in the unrelated genera *Hemidactylus* and *Luperosaurus* (Dudley et al., 2007; Heinicke et al., 2012).

### 4.3. Classification

Given our resolution of three major clades of *Gekko*—one comprising almost exclusively Philippine species, another primarily Southeast Asian mainland (plus Sundaland) taxa, and a third consisting of morphologically variable species from Southeast Asia, Wallacea, Melanesia and the Southwest Pacific—we subsume all contained taxa into the oldest, most inclusive generic name, *Gekko* (Laurenti). This solution imparts the fewest alterations of binominal species named pairs, and thus, is the most conservative and preferred option. Accordingly, we place all species of the genus *Ptychozoon*, as well as the Sulawesi endemics *Luperosaurus iskandari* and *L. browni*, and the Palawan *L. gulat*, into the genus *Gekko*. In consideration of *Gekko* phylogenetic relationships (Rösler et al., 2011; Brown et al., 2012a; Heinicke et al., 2012; this study), distinctly geographically circumscribed regional clades (Japan and adjacent mainland, Indochina, southwest Pacific, the Philippine archipelago), plus associated morphological variability of the contained taxa, we find the recognition of the following six phylogenetically and phenotypically defined subgenera advisable at this time.

Genus *Gekko* (Fig. 3, clades A–G)

Subgenus *Gekko* (Fig. 3A)

Type species: *Gekko*(*Gekko*) *gecko* (Linnaeus, 1758), designated by tautonymy fide Stejneger (1936).

Definition: Subgenus *Gekko* is a maximum crown-clade name referring to the clade originating with the most recent common ancestor of *Gekko* (*Gekko*) *gecko*, and *G*. (*G*.) *smithii*, and including all extant species that share a more recent common ancestor with these taxa than with any of the type species of other sub-genera recognized here. Although unambiguous synapomorphies for this group have not been identified, members of the subgenus *Gekko* are larger than most conspecifics (adults SVL > 110 mm), with tubercles present on ventrolateral folds, more than 18 subdigital Toe IV scansors, femoral pores absent, and with a relatively low number of precloacal pores (Bauer et al., 2008).

Content: *Gekko* (*Gekko*) *G. albofasciolatus* Günther, 1867, *G*. (*G*.) *gecko* (Linnaeus, 1758), *G*. (*G*.) *nutaphandi* BauerSumontha & Pauwels, 2008; *G*. (*G*.) *reevesii*(Gray, 1831), *G*. (*G*.) *siamensis* Grossmann & Ulber, 1990, *G*. (*G*.) *smithii* Gray, 1842, *G*. (*G*.) *verreauxi* Tytler, 1864.

Comment: The subgenus includes taxa from Rösler et al. (2011) *G. gecko* Group. In recognizing this assemblage, Rösler et al. (2011) resurrected *G. reevesii* and provided data on two subspecies of *G*. (*G*.) *gecko*. Undescribed species in this subgenus have been reported from Sulawesi, the Togian Islands of Indonesia, and possibly the islands of Tioman and Tulai of Malaysia (Grismer, 2006; Rösler et al., 2011).

Subgenus *Japonigekko* subgen. nov. (Fig. 3B)

Type species: *Gekko* (*Japonigekko*) *japonicus* (Schlegel, 1836), the oldest name available for a taxon included in the new subgenus, here designated.

Definition: *Gekko* (Subgenus *Japonigekko*) is a maximum crown-clade name referring to the clade originating with the most recent common ancestor of *Gekko* (*Japonigekko*) *chinensis*, *G*. (*J*.) *swinhonis*, and including all extant species that share a more recent common ancestor with these species than with any of the type species of the other subgenera recognized here. Members of the subgenus *Japonigekko* are usually relatively small to moderate-sized taxa (59–99 mm SVL); possess or lack dorsal tubercle rows (a few species possess up to 21 rows); possess up to 32 precloacal pores (most lack pores altogether); and lack tubercles on ventrolateral folds. All contained taxa possess some degree of interdigital webbing (minimally to extensively webbed). Other than the presence of interdigital webbing, unambiguous synapomorphies for the new subgenus have not been identified; nevertheless, our phylogeny strongly corroborates the monophyly of this phenotypically variable clade.

Content: *Gekko* (*Japonigekko*) *adleri* Nguyen, Wang, Yang, Lehmann, Le, Ziegler and Bonkowski, 2013; *G*. (*J*.) *aaronbaueri* Tri, Thai, Phimvohan, David, and Teynié, 2015; *G. (*J*.) *auriverrucosus* Zhou & Liu, 1982; *G*. (*J*.) *bonkowskii* Luu, Calme, Nguyen, Le, and Ziegler, 2015; *G*. (*J*.) *canhi* Rösler, Nguyen, Doan, Ho & Ziegler, 2010; *G*. (*J*.) chinensis* Gray, 1842; *G*. (*J*.) *guishanicus* Lin & Yao, 2016; *G*. (*J*.) *hokouensis* Pope, 1928; *G*. (*J*.) *japonicus* (Schlegel, 1836); *G*. (*J*.) *kwangsiensis* Yang, 2015; *G*. (*J*.) *lauhachindai* Yang 2015; *G*. (*J*.) *liboensis* Zaho & Li, 1982; *G*. (*J*.) *melli* Vogt, 1922; *G*. (*J*.) *nadenensis* Luu, Nguyen, Le, Bonkowski, Ziegler, 2017; *G*. (*J*.) *palmatus* Boulenger, 1907; *G*. (*J*.) *scabridus* Liu & Zhou, 1982; *G*. (*J*.) *scientiadventura* Röosler, Ziegler, Vu, Herrmann & Böhme, 2004; *G*. (*J*.) *sengchanthavongi* Luu, Calme, Nguyen, Le, and Ziegler 2015; *G*. (*J*.) *shibatai* Toda, Sengoku, Hikida & Ota, 2008; *G*. (*J*.) *similignum* Smith, 1923; *G*. (*J*.) *subpalmatus* Günther, 1864; *G*. (*J*.) *swinhonis* Günther, 1864; *G*. (*J*.) *taibaiensis* Song, 1985; *G*. (*J*.) *tawaensis* Okada, 1956; *G*. (*J*.) *thakhekensis* Luu, Calme, Nguyen, Le, Bronkowski, and Ziegler, 2014; *G*. (*J*.) *truongi* Phung and Ziegler, 2011; *G*. (*J*.) *vertebralis* Toda, Sengoku, Hikida & Ota, 2008; *G*. (*J*.) *vietnamensis* Sang, 2010 *incertae sedis G*. (*J*.) *wenxianensis* Zhou and Wang, 2008; *G*. (*J*.) *yakuensi* Matsui & Okada, 1968.

Comment: The subgenus includes taxa from the *G. japonicus* Group of Rösler et al. (2011). See Rösler et al. (2011) for discussions of the considerable interspecific morphological variability and lengthy taxonomic controversy that characterizes this assemblage (Boulenger, 1885, 1907; Smith, 1935; Bourret, 1937; Bauer, 1994; Ota et al., 1995; Nguyen et al., 2009).

Etymology: *Japonigekko* is a masculine noun, referring to geographic origin (Japan) of the type species (*G. japonicus)*. We use the spelling “Japon” following Schlegel (1836) who used this French spelling in the construction of the name *Platydactylus japonicus*.

Subgenus *Ptychozoon* (Fig. 3C)

Type species: *Ptychozoon homalocephala* (Creveldt, 1809) (preoccupied [as *Lacerta homalocephala*]; nomen novum *P. kuhli* proposed by Stejneger, 1902), the oldest name available for a taxon included in this subgenus, here designated.

Definition: *Gekko* (Subgenus *Ptychozoon*) is a maximum crown-clade name referring to the clade originating with the most recent common ancestor of *Ptychozoon horsfieldii* (Gray, 1827) *Gekko* (*Ptychozoon*) *kuhli*, and including all extant species that share a more recent common ancestor with these taxa than with any of the type species of other subgenera recognized here. Unambiguous synapomorphies for the subgenus *Ptychozoon* include the presence of a midbody axilla–groin patagial membrane (“parachute”), denticulate dermal lobes along lateral margins of the tail, expanded dermal flaps on anterior and posterior margins of the limbs, and the presence of extensive interdigital webbing of the hands and feet. Additionally, all taxa (the first-branching *Ptychozoon* lineage [Brown et al., 2012a, b]) possess enlarged, imbricate parachute support scales, an infraauricular cutaneous flap (“canard wing”), and a variably enlarged terminal tail flap.

Content: *Gekko* (*Ptychozoon*) *banannensis* Wang, Wang, and Liu, 2016, *G*. (*P*) *cicakterbang* Grismer, Wood, Grismer, Quah, Thy, Phimmachak, Sivongxay, Seateun, Stuart, Siler, Mulcahy, and Brown, 2019, *G*. (*P*.) *horsfieldii* (Gray, 1827), *G*. (*P*.) *intermedium* Taylor 1915, *G*. (*P*.) *kabkaebin* Grismer, Wood, Grismer, Quah, Thy, Phimmachak, Sivongxay, Seateun, Stuart, Siler, Mulcahy, and Brown, 2019, *G*. (*P*.) *kaengkrachanense* Sumontha, Pauwels, Kunya, Limlikhitaksorn, Ruksue, Taokratok, Ansermet, and Chanhome, 2012, *G*. (*P*.) *kuhli*(Stejnegeri, 1902). *G*. (*P*.) *lionotum* Annandale, 1905, *G*. (*P*.) *tokehos* (Grismer et al. 2019), *G*. (*P*.) *nicobarensis* Das and Vijayakumar 2009, *G*. (*P*.) *popaense* Grismer, Wood, Thura, Grismer, Brown, Stuart, 2018, *G*. (*P*.) *tokehos* Grismer, Wood, Grismer, Quah, Thy, Phimmachak, Sivongxay, Seateun, Stuart, Siler, Mulcahy, and Brown, 2019, and *G*. (*P*.) *trinotaterra* Brown, 1999.

Comment: In revising species “group” classification of *Gekko*, Rösler et al. (2011) did not comment on the inclusion of *Ptychozoon* species (unsampled in that study), which were later found to be imbedded within *Gekko* (Brown et al., 2012a, b; Heinicke et al., 2012).

Subgenus *Rhacogekko* subgen. nov. (Fig. 3D)

Type species: *Gekko* (*Rhacogekko*) *rhacophorus* (Boulenger, 1899), the oldest (and only) name available for a taxon included in the new subgenus, here designated.

Definition: *Rhacogekko* (new Subgenus) is a maximum crown-lineage name referring to *Gekko* (*R*.) *rhacophorus* originating as the sister lineage to *Sundagekko* and *Pseudosundagekko. Gekko* (*R*.) *rhacophorus*. Although unambiguous synapomorphies for the new subgenus have not been identified, *Gekko* (*R*.) *rhacophorus* differs from other subgenera by lacking lateral skin folds on head (infra-auricular cutaneous flaps), lacking imbricate dorsal parachute support scales and an enlarged terminal tail flap (Brown et al., 1997), and by having fully half-webbed fingers and toes (Boulenger, 1899; Malkamus et al., 2002).

Content: *Gekko* (*R*.) *rhacophorus* (Boulenger, 1899) and *Gekko* (*R*.) *sorok*(Das et al. 2008) *incertae sedis*.

Etymology: *Rhacogekko* is derived from the Greek noun *rhakos*, meaning “rag” or “wrinkle” in relation to the rounded lobe-like fringes or wrinkles on the lateral folds of the body.

Subgenus *Sundagekko* subgen. nov. (Fig. 3E)

Type species: *Gekko* (*Sundagekko*) *vittatus* Houttuyn, 1782, the oldest name available for a taxon included in the new subgenus, here designated.

Definition: *Sundagekko*, new subgenus, is a maximum crown-clade name referring to the clade originating with the most recent common ancestor of *Gekko* (*Sundagekko*) *vittatus, G*. (*P*.) *browni*, and *G*. (*P*.) *iskandari* (new combination) and including all extant species that share a more recent common ancestor with these taxa than with any of the type species of other subgenera recognized here. Unambiguous synapomorphies for the new subgenus have not been identified, but three known (and two inferred) members of *Sundagekko* are gracile, slender taxa (*G*. [*S*.] *iskandari, G*. [*S*.] *vittatus, G*. [*S*.] *browni*, presumably *G*. [*S*.] *brooksi* and *G*. [*S*.] *remotus*) with thin, elongate bodies and ornate tubercles present on ventrolateral folds (Brown et al., 2000; McCoy, 2006; Rösler et al., 2012).

Content: *Gekko* (*Sundagekko) brooksi*, Boulenger, 1920 (new combination), *Gekko* (*S*.) *browni*, (Russell, 1979), *Gekko* (*S*.) *flavimaritus* Rujirawana, Fong, Ampai, Yodthong, Termprayoon, and Anchalee Aowphol, 2019; *Gekko* (*S*.) *iskandari*, (Brown, Supriatna, and Ota, 2000) (new combination), *G*. (*S*.) *remotus* Rösler, Ineich, Wilms, and Böhme, 2012; and *G*. (*S*.) *vittatus* Houttuyn, 1782; *Gekko* (*Petrigekko*) *badenii* Nekrasova & Szczerbak, 1993; *G*. (*S*.) *boehmei* Luu, Calme, Nguyen, Le, and Ziegler, 2015; *G*. (*S*.) *canaensis* Ngo & Gamble, 2011; *G*. (*S*.) *grossmanni* Günther, 1994; *G*. (*S*.) *lauhachindaei* Panitvong, Sumontha, Konlek & Kunya, 2010; *G*. (*S*.) *petricolus* Taylor, 1962; *G*. (*S*.) *russelltraini* Ngo, Bauer, Wood & Grismer, 2009; *G*. (*S*.) *takouensis* Ngo & Gamble, 2010.

Comment: Our phylogenomic approach confirms the phenotypically dissimilar pairing of the taxa *Gekko* (*S*.) *vittatus* (plus the closely-related *G. remotus*) and *G*. (*S*.) *iskandari* (inferred previously with two genes; (Brown et al., 2012a). Despite their morphological differences (Crombie and Pregill, 1999; Brown et al., 2000; McCoy, 2006), we note that the proximate geographic ranges in the islands of Melanesia, Palau, and the southwest Pacific lend geographical support to the recognition of the new subgenus (Brown et al., 2000; McCoy, 2006).

Etymology: *Sundagekko* is a masculine noun, derived from the English noun formed from the combination of the term “Sunda” and “gekko,” collectively referring to the continental extension of Indochina referred to as Sundaland, which is composed of the Malay Peninsula, Sumatra, Borneo, Java, Bali and other small Island archipelagos in the Western Philippine Sea. Although *G*. (*S*.) *iskandari, G*. (*S*.) *vittatus*, and *G*. (*S*.) *remotus* are distributed beyond the confines of the Sunda Shelf, the species *G*. (*P*.) *brooksi* (Sumatra Island), and *G*. (*P*.) *browni* (Malaysian Peninsula, Borneo) are known only within Sundaland.

Subgenus *Pseudosundagekko* subgen. nov. (Fig. 3F)

Type species: *Gekko*(*P*.) *gulat* (Brown, Diesmos, Duya, Garcia, and Rico, 2010), here designated.

Definition: *Pseudosundagekko* is the sister lineage to a crown clade of Philippine endemics described below. At present containing a single species (*G*.[*P*.] *gulat*), the new subgenus is intended to include any species discovered in the future to share a more recent common ancestor with *G*.(*P*.) *gulat* than with any of the type species of other subgenera recognized here.

Content: *Gekko* (*P*.) *gulat* (Brown, Diesmos, Duya, Garcia, and Rico, 2010).

Etymology: *Pseudosundagekko* is a masculine noun, formed in reference “Pseudo,” meaning false, “Sunda” in reference to the landmasses of the Sunda Shelf, and “gecko,” for gekkonid lizards; this name is chosen in reference to recent changes in understanding biogeographical relationships and evolutionary history of many iconic vertebrate lineages endemic to the island of Palawan Island (Blackburn et al., 2010; Esselstyn et al., 2010; Siler et al., 2012; Brown et al., 2016), erroneously and over simplistically assumed previously to be a simple faunal extension of the Sunda Shelf island of Borneo.

Subgenus *Archipelagekko* subgen. nov. (Fig. 3G)

Type species: *Gekko* (*Archipelagekko*) *mindorensis* Taylor, 1919, the oldest name available for a taxon included in the new subgenus.

Definition: *Archipelagekko* is a maximum crown-clade name referring to the clade originating with the most recent common ancestor of *Gekko* (*Archipelagekko*) *athymus* and includes all extant species that share a more recent common ancestor with these taxa than with any of the type species of other subgenera recognized here.

Content: *Gekko* (*Archipelagekko*) *athymus* Brown and Alcala, 1962, *G*. (*A*.) *carusadensis* Linkem, Siler, Diesmos, Sy, and Brown, 2010, *G*. (*A*.) *coi* Brown, Siler, Oliveros, and Alcala, 2011, *G*. (*A*.) *crombota* Brown, Oliveros, Siler, and Diesmos, 2006, *G*. (*A*.) *ernstkelleri* Rösler, Siler, Brown, Demeglio, and Gaulke, 2006, *G*. (*A*.) *gigante* Brown and Alcala 1978, *G*. (*A*.) *kikuchii* (Oshima, 1912), *G*. (*A*.) *monarchus* (Schlegel, 1836), *G*. (*A*.) *mindorensis* Taylorm, 1919, *G*. (*A*.) *palawanensis* Taylor, 1925, *G*. (*A*.) *romblon* Brown and Alcala 1978, and *G*. (*A*.) *rossi* Brown, Oliveros, Siler, and Diesmos, 2009.

Comment: The new subgenus contains *G. athymus* (not previously placed in a species group due to its morphological distinctiveness; Rösler et al., 2011), the *G. porosus* Group, and the *G. monarchus* Group (Rösler et al., 2011). We are aware of at least two additional unrecognized species in this clade (G. sp. A. [Dalupiri Isl.] and *G*. sp. B [Camiguin Norte Isl.]; Brown et al. 2009); all are restricted to the Philippines, with the exception of *G. monarchus* (a widespread taxon that is known from Palawan [Philippines], parts of Indonesia, Malaysia, and Thailand [Rösler et al., 2011; Grismer, 2011; Siler et al., 2012], and *G. kikuchii*, a northern Philippine species with a distribution extending to Lanyu Island, Taiwan [Oshima, 1912; Siler et al., 2014]). An additional 4–7 species likely currently reside within the synonymy of *G*. (*A*.) *mindorensis* (Siler et al., 2014).

Etymology: *Archipelagekko* is a masculine noun, derived from the English noun Archipelago, in recognition of the observation that nearly all contained taxa are restricted to the Philippine Archipelago. Suggested common name: Philippine Geckos.

### 4.4. Taxonomic vandalism

While this manuscript was in preparation, a series of three papers appeared (Hoser, 2018a, b, c,) in which revised taxonomies were presented for *Gekko, Luperosaurus*, and *Ptychozoon*, as well as *Lepidodactylus* and *Pseudogekko*. The basis for these papers were the various publications cited herein (e.g., Rösler et al., 2011; Brown et al., 2012a, b; Heinicke, 2012; Oliver et al., 2018) in which phylogenetic hypotheses based on Sanger sequencing results were presented, without concomitant taxonomic rearrangements, pending the collection of data which would permit more advanced analyses, and well-supported topologies (this work). Hoser (2018a, b, c) proposed several new generic and subgeneric names to deal with cases of paraphyly and polyphyly as they were understood and misunderstood by him, prior to the present study. Additionally, he proposed numerous new species names, based on unsubstantiated (i.e., presented without accompanying data from voucher specimens) and operationally non-diagnostic “diagnoses” based, apparently, on “character states” gleaned from figures/photographs included in Brown et al. (2012a, b), Heinicke et al. (2012), and Oliver et al. (2018). Aside from his disregarding of conventions of species descriptions, some of his higher taxonomic proposals are partly, though not entirely, congruent with the units recognized herein and also include some generic proposals that our current data suggest are non-monophyletic. Although acknowledging the temporal precedence of Hoser’s names, we follow Kaiser et al. (2013) and Kaiser (2014) in rejecting Hoser’s taxonomic vandalism and, for the reasons cited therein, we regard these names as unavailable for nomenclatural purposes. We echo the belief that the extraordinary chaos and rampant ethical violations resulting from Hoser’s actions require urgent use of the plenary powers by the ICZN. Without such action, legitimate scientists must knowingly violate the principle of priority or risk that their own original research be used to fuel the ongoing deluge of taxonomic vandalism (Borrell, 2007).

If widely adopted during the standard process of phylogeny-based revisionary classification, the practice of formally proposing available subgeneric names to recognize phylogenetically identified and defined clades and lineages has an added practical benefit of potentially alleviating the egocentric, looting motivation behind taxonomic vandalism. Whether defined by standard Linnaean system, character-based diagnoses (Vences and Glaw, 2006), phylogenetically-defined and rank free (Hillis and Wilcox, 2005, Leaché et al., 2009; Wallach et al., 2009; Yuan et al., 2016), or with parallel classifications that embrace both systems (Brown et al., 2013), the existence of a node-based subgenus name (defined with reference to a phylogeny) in synonymy with a more inclusive genus name, and satisfying the qualifying conditions of The Code (including a character-based diagnosis and designation of a type species), alleviates the threat of taxonomic vandalism via the simple requirement that the subgenus name be used should future systematists effectively argue for a more atomized classification. Although this simple practice may involve additional behind-the-scenes steps (defining and naming supraspecific taxa that will not be used in today’s binomial classification), it effectively eliminates the nefarious motivation behind taxonomic vandalism, while ensuring that supraspecific names are created only when accompanied by phylogenetic data, and that they explicitly involve appropriate specifiers in the form of type species and, ultimately, type specimens.

## Acknowledgments

Funding for UCE sequence capture work at KU was provided (to RMB and RGM) by the University of Kansas Office of the Provost, Research Investment Council; Illumina sequencing performed at the KU Genome Sequencing Core facility, and Southeast Asian fieldwork was supported by grants from the U.S. National Science Foundation (NSF): DEB 0743491, 1418895, 0640737, 0344430, 0073199, 1654388 and EF-0334952 to RMB; and DEB 0804115 and 1657648 and IOS 1353683 to CDS. XG was supported by grants from the Study Abroad Scholarship Program of the Chinese Academy of Sciences, and the National Natural Science Foundation of China (31672270). RGM was supported by NSF award DEB-1557053. AMB was supported by NSF grants DEB 0844523, 1555968 and EF 1241885 (subaward 13-0632) and by the Gerald M. Lemole Endowed Chair Funds. We would like to thank Evan S.H. Quah and Kurt H.P. Guek for providing images of *G*. (*J*.) *japonicus* and *G*. (*R*.) *rhacophorus* for the graphical abstract. This paper is contribution number XXX of the Auburn University Museum of Natural History.

---

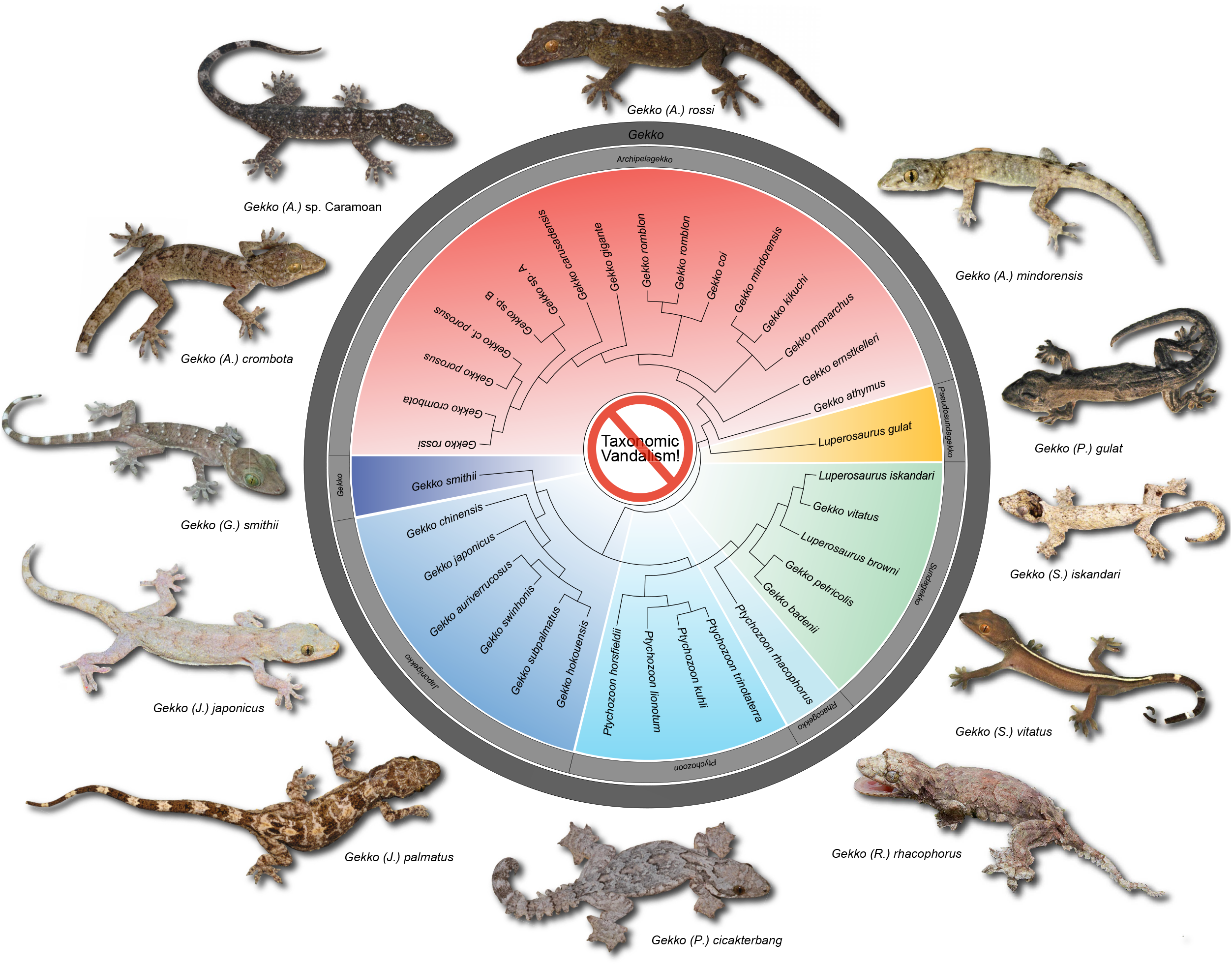

## References

Barbour, T., 1912. A contribution to the zoogeography of the East Indian Islands. Mem. Mus. Comp. Zool. 44, 1–203.

Bankevich, A., Nurk, S., Antipov, D., Gurevich, A.A., Dvorkin, M., Kulikov, A.S., Lesin, V.M., Nikolenko, S.I., Pham, S., Prjibelski, A.D., 2012. SPAdes: a new genome assembly algorithm and its applications to single-cell sequencing. J. Comput. Biol. 19, 455–477.

Bayzid, M.S., Warnow, T., 2012. Estimating optimal species trees from incomplete gene trees under deep coalescence. J. Comput. Biol. 19, 591–605.

Bauer, A.M., 1994. Familia Gekkonidae (Reptilia, Sauria). Part I: Australia and Oceania. Walter de Gruyter, Berlin.

Bauer, A.M., Sumontha, M., Pauwels, O.S., 2008. A new red-eyed *Gekko* (Reptilia: Gekkonidae) from Kanchanaburi province, Thailand. Zootaxa 1750, 32–42.

Blackburn, D.C., Bickford, D.P., Diesmos, A.C., Iskandar, D.T., Brown, R.M., 2010. An ancient origin for the enigmatic Flat-Headed Frogs (Bombinatoridae: *Barbourula*) from the islands of Southeast Asia. PLOS ONE 5, e12090.

Borrell, B. 2007. Linnaeus at 300: The big name hunters. Nature 446: 253–255.

Boulenger, G.A., 1907. Descriptions of new lizards in the British museum. Ann. Mag. Nat. Hist. 19, 486–489.

Boulenger, G.A., 1885. Catalogue of the Lizards in the British Museum volume 2. Trustees of the British Museum, London.

Boulenger, G.A., 1899. LI.—Descriptions of three new reptiles and a new Batrachian from Mount Kina Balu, North Borneo. J. Nat. Hist. 4, 451–454.

Bourret, R., 1937. Notes herpétologiques sur l’Indochine française. XV. Lézards et serpents reçu au laboratoire des Sciences Naturelles de l’Université au cours de l’année 1937. Descriptions de deux espèces et de deux variétés nouvelles. Bulletin Générale de l’Instruction Publique 5, 57–82.

Brown, R.M., Diesmos, A.C., Duya, M.V., Garcia, H.J.D., Rico, E.L., 2010. A new forest gecko (Squamata; Gekkonidae; genus *Luperosaurus*) from Mt. Mantalingajan, southern Palawan Island, Philippines. J. Herpetol. 44, 37–48.

Brown, R.M., Diesmos, A.C., Duya, M.V., 2007. A new species of *Luperosaurus* (Squamata: Gekkonidae) from the Sierra Madre mountain range of northern Luzon Island, Philippines. Raffles Bull. Zool. 55, 153–160.

Brown, R.M., Ferner, J.W., Diesmos A.C., 1997. Definition of the Philippine parachute gecko, *Ptychozoon intermedium* Taylor 1915 (Reptilia: Squamata: Gekkonidae): redescription, designation of a neotype, and comparisons with related species. Herpetologica 53, 357–373.

Brown, R.M., Oliveros, C., Siler, C.D., Diesmos, A.C., 2009. Phylogeny of *Gekko* from the northern Philippines, and description of a new species from Calayan Island. J. Herpetol. 43, 620–635.

Brown, R.M., Siler, C.D., Das, I., Min, Y., 2012a. Testing the phylogenetic affinities of Southeast Asia’s rarest geckos: flap-legged geckos (*Luperosaurus*), flying geckos (*Ptychozoon*) and their relationship to the pan-Asian genus *Gekko*. Mol. Phylogenet. Evol. 63, 915–921.

Brown, R.M., Siler, C.D., Grismer, L.L., Das, I., McGuire, J.A., 2012b. Phylogeny and cryptic diversification in southeast Asian flying geckos. Mol. Phylogenet. Evol. 65, 351–361.

Brown, R.M., Supriatna, J., Ota, H., 2000. Discovery of a new species of *Luperosaurus* (Squamata; Gekkonidae) from Sulawesi, with a phylogenetic analysis of the genus, and comments on the status of *Luperosaurus serraticaudus*. Copeia 2000, 191–209.

Brown, R.M., Siler, C.D., Oliveros, C.H., Esselstyn, J.A., Diesmos, A.C., Hosner, P.A., Linkem, C.W., Barley, A.J., Oaks, J.R., Sanguila, M.B., Welton, L.J., Blackburn, D.S., Moyle, R.G., Peterson, A.T., Alcala, A.C., 2013. Evolutionary processes of diversification in a model island archipelago. Annu. Rev. Ecol. Evol. Syst. 44, 411–435.

Brown, R.M., Su, Y.-C., Barger, B., Siler, C.D., Sanguila, M.B., Diesmos, A.C., Blackburn, D. C., 2016. Phylogeny of the island archipelago frog genus *Sanguirana:* another endemic Philippine radiation that diversified ‘Out-of-Palawan’. Mol. Phylogenet. Evol. 94, 531–536.

Capella-Gutiérrez, S., Silla-Martínez, J. M., Gabaldón, T., 2009. trimAI: a tool for automated alignment trimming in large-scale phylogenetic analyses. Bioinformatics 25, 1972–1973.

Castillo-Ramírez, S., Liu, L., Pearl, D., Edwards, S.V., 2010. Bayesian estimation of species trees: a practical guide to optimal sampling and analysis. In: Knowles, L.L, Kubatko, L.S. (Eds.), Estimating Species Trees: Practical and Theoretical Aspects, Hoboken: Wiley-Blackwell. Pp. 15–33.

Chan, K.O., Brown, R.M., 2017. Did true frogs ‘dispersify’? Biol. Lett. 13, 20170299.

Chen, S., Huang, T., Zhou, Y., Han, Y., Xu, M., Gu, J., 2017. AfterQC: automatic filtering, trimming, error removing and quality control for fastq data. BMC Bioinformatics 18, 80.

Chifman, J., Kubatko, L., 2014. Quartet inference from SNP data under the coalescent model. Bioinformatics 30, 3317–3324.

Chou, J., Gupta, A., Yaduvanshi, S., Davidson, R., Nute, M., Mirarab, S., Warnow, T., 2015. A comparative study of SVDquartets and other coalescent-based species tree estimation methods. BMC Genomics 16, S2.

Crombie, R.I., Pregill, G.K., 1999. A checklist of the herpetofauna of the Palau Islands (Republic of Belau), Oceania. Herpetol. Monogr. 13, 29–80.

Dudley, R., Byrnes, G., Yanoviak, S., Borrell, B., Brown, R.M., McGuire, J., 2007. Gliding and the functional origins of flight: biomechanical novelty or necessity? Annu. Rev. Ecol. Evol. Syst. 38, 179–201.

Edwards, S.V., Liu, L, Pearl, D.K., 2007. High-resolution species trees without concatenation. Proc. Natl. Acad. Sci. USA 104, 5936–5941.

Edwards, S.V., Xi, Z., Janke, A., Faircloth, B.C., McCormack, J.E., Glenn, T.C., Zhong, B., Wu, S., Lemmon, E.M., Lemmon, A.R., Leaché, A.D., Liu, L., Davis, C.C., 2016. Implementing and testing the multispecies coalescent model: a valuable paradigm for phylogenomics. Mol. Phylogenet. Evol. 94, 447–462.

Esselstyn, J.A., Oliveros, C.H. Moyle, R.G., Peterson, A.T., McGuire, J.A., Brown, R.M., 2010. Integrating phylogenetic and taxonomic evidence illuminates complex biogeographic patterns along Huxley’s modification of Wallace’s Line. J. Biogeogr. 37, 2054–2066.

Faircloth, B., McCormack, J., Crawford, N., Harvey, M., Brumfield, R., Glenn, T., 2012. Ultraconserved elements anchor thousands of genetic markers spanning multiple evolutionary timescales. Sys. Biol. 61, 717–726.

Gamble, T., Greenbaum, E., Jackman, T.R., Russell, A.P., Bauer, A.M., 2012. Repeated origin and loss of adhesive toepads in geckos. PLoS One 7, e39429.

Glaw, F., & Vences, M. (2006). Phylogeny and genus-level classification of mantellid frogs (Amphibia, Anura). Organisms Diversity & Evolution, 6(3), 236–253.

Glenn, T.C., Nilsen, R., Kieran, T.J., Finger, J.W., Pierson, T.W., Bentley, K.E., Hoffberg, S., Louha, S., Garcia-De-Leon, F.J., Angel del Rio Portilla, M., Reed, K., Anderson, J.L., Meece, J.K., Aggery, S., Rekaya, R., Alabady, M., Belanger, M., Winker, K., Faircloth, B.C., 2016. Adapterama I: universal stubs and primers for thousands of dual-indexed illumina libraries (itru & inext). bioRxiv. URL: https://www.biorxiv.org/content/early/2016/06/15/049114. doi:10.1101/049114. arXiv:https://www.biorxiv.org/content/early/2016/06/15/049114.full.pdf.

Grismer, L.L., 2006. Amphibians and reptiles of the Tioman Archipelago, Malaysia. Forestry Department Peninsular Malaysia, Kuala Lumpur.

Grismer, L.L., 2011. Field Guide to the Amphibians and Reptiles of the Seribuat Archipelago (Peninsular Malaysia). Edition Chimaira, Frankfurt am Main.

Grismer, L.L., Wood, Jr, P.L., Aowphol, A., Cota, M., Grismer, M.S., Murdoch, M.L., Aguilar, C., Grismer, J.L., 2016. Out of Borneo, again and again: biogeography of the Stream Toad genus *Ansonia* Stoliczka (Anura: Bufonidae) and the discovery of the first limestone cave-dwelling species. Biol. J. Linnean. Soc. 120, 371–395.

Grismer, L.L., Wood, Jr., P.L., Thura, M.K., Grismer, M.S., Brown, R.M., Stuart, B.L. 2018. Geographically structured genetic variation in *Ptychozoon lionotum* (Squamata: Gekkonidae) and a new species from an isolated volcano in Myanmar. Zootaxa 4514, 202–214.

Grismer, L.L., Wood, Jr., P.L., Grismer, J.L., Quah, E.S.H., Thy, N., Phimmachak, S., Sivongxay, N., Seateun, S., Stuart, B.L., Siler, C.D., Mulcahy, D.G., Anamza, T., Brown, R.M. Phylogeographic structure of the Parachute Gecko *Ptychozoon lionotum* Annandale, 1905 across Indochina and Sundaland with descriptions of three new species. Zootaxa 4638, 151–198.

Heinicke, M.P., Greenbaum, E., Jackman, T.R., Bauer, A.M., 2012. Evolution of gliding in Southeast Asian geckos and other vertebrates is temporally congruent with dipterocarp forest development. Biol. Lett. 8, 994–997.

Heyer, W.R., Pongsapipatana, S., 1970. Gliding speeds of *Ptychozoon lionotum* (Reptilia: Gekkonidae) and *Chrysopelea ornata* (Reptilia: Colubridae). Herpetologica 26, 317–319.

Hillis, D.M., Wilcox, T.P., 2005. Phylogeny of the New World true frogs (Rana). Mol. Phylogenet. Evol. 34, 299–314.

Hoser, R.T., 2018a. A significant improvement to the taxonomy of the gecko genus *Gekko* Laurenti, 1768 *sensu lato* to better reflect morphological diversity and ancient divergence within the group. Australasian J. Herpetol. 38, 6–18.

Hoser, R.T., 2018b. A revised taxonomy of the gecko genus *Ptychozoon* Kuhl and Van Hasselt, 1822, including the formal erection of two new genera to accommodate the most divergent taxa and description of ten new species. Australasian J. Herpetol. 38, 19–31.

Hoser, R.T., 2018c. A revised taxonomy of the gecko genera *Lepidodactylus* Fitzinger, 1843, *Luperosaurus* Gray, 1845 and *Pseudogekko* Taylor, 1922 including the formal erection of new genera and subgenera to accommodate the most divergent taxa and description of 26 new species. Australasian J. Herpetol. 38, 32–64.

Kaiser, H., 2014. Best practices in herpetological taxonomy: errata and addenda. Herpetol. Rev. 45, 257–268.

Kaiser, H., Crother, B.I., Kelly, C.M., Luiselli, L., O’Shea, M., Ota, H., Passos, P., Schleip, W.D. Wüster, W., 2013. Best practices: in the 21st century, taxonomic decisions in herpetology are acceptable only when supported by a body of evidence and published via peer-review. Herpetol. Rev. 44, 8–23.

Kalyaanamoorthy, S., Minh, B.Q., Wong, T.K., von Haeseler, A. Jermiin, L.S., 2017. Modelfinder: fast model selection for accurate phylogenetic estimates. Nat. Methods 14, 587.

Katoh, K., Standley, D., 2013. MAFFT multiple sequence alignment software version 7: improvements in performance and usability. Mol. Biol. Evol. 30, 772–80.

Kent, W.J., 2002. Blat—the blast-like alignment tool. Genome Res. 12, 656–664.

Kubatko, L., Degnan, J., 2007. Inconsistency of phylogenetic estimates from concatenated data under coalescence. Syst. Biol. 56, 17–24.

Leaché, A.D., Koo, M.S., Spencer, C.L., Papenfuss, T.J., Fisher, R.N., McGuire, J.A., 2009. Quantifying ecological, morphological, and genetic variation to delimit species in the coast horned lizard species complex (*Phrynosoma*). Proc. Natl. Acad. Sci. USA 106(30), 12418–12423.

Malkmus, R, Manthey, U., Vogel, G., Hoffman, P., Kosuch, J., 2002. Amphibians and Reptiles of Mount Kinabalu (North Borneo). Ganter Verlag, Ruggell, Germany.

Marcellini, D.L., Keefer, T.E., 1976. Analysis of the gliding behavior of *Ptychozoon lionatum* (Reptilia: Gekkonidae). Herpetologica 32, 362–66.

McCoy, M., 2006. Reptiles of the Solomon Islands. Pensoft.

Meng, W., 2018. pblat – blat with multi-threads support. URL: http://icebert.github.io/pblat/.

Minh, Q., Nguyen, M., von Haeseler, A.A., 2013. Ultrafast approximation for phylogenetic bootstrap. Mol. Biol. Evol. 30, 1188–1195.

Nguyen, V., Ho, T., Nguyen, Q., 2009. Herpetofauna of Vietnam. Edition Chimaira, Frankfurt am Main, Germany.

Nguyen, L.-T., Schmidt, H.A., von Haeseler, A., Minh, B.Q., 2015. IQ-TREE: a fast and effective stochastic algorithm for estimating maximum-likelihood phylogenies. Mol. Biol. Evol. 32, 268–274.

Oliver, P., Brown, R.M., Kraus, F., Rittmeyer, E., Travers, S.L., Siler, C.D., 2018. Lizards of the lost arcs: mid-Cenozoic diversification, persistence and ecological marginalization in the West Pacific. Proc. Biol. Sci. B. 285, 20171760.

Oshima, M., 1912. Description of a new gecko from Botel Tobago island. Philippine J. Sci. 7, 241–242.

Ota, H., Lau, M., Weidenhöfer, T., Yasukawa, Y., Bogadek, A., 1995. Taxonomic review of the geckos allied to *Gekko chinensis* Gray 1842 (Gekkonidae Reptilia) from China and Vietnam. Trop. Zool. 8, 181–196.

Pyron, R.A., Burbrink, F.T. Wiens, J.J., 2013. A phylogeny and revised classification of Squamata, including 4161 species of lizards and snakes. BMC Evol. Biol. 13, 93.

Rösler, H., Bauer, A.M., Heinicke, M.P., Greenbaum, E., Jackman, T., Nguyen, T.Q., Ziegler, T., 2011. Phylogeny, taxonomy, and zoogeography of the genus *Gekko laurenti*, 1768 with the revalidation of *G. reevesii* Gray, 1831 (Sauria: Gekkonidae). Zootaxa 2989, 1–50.

Rösler, H., Ineich, I., Wilms, T.M., Böhme, W., 2012. Studies on the taxonomy of the *Gekko vittatus* Houttuyn, 1782 complex (Squamata: Gekkonidae) I. On the variability of *G. vittatus* Houttuyn, 1782 sensu lato, with the description of a new species from Palau Islands, Micronesia. Bonn Zool. Bull. 61, 241–254.

Russell, A., 1979. The origin of parachuting locomotion in gekkonid lizards (Reptilia: Gekkonidae). Zool. J. Linn. Soc. 65, 233–249.

Russell, A.P., Dijkstra, L.D., Powell, G.L., 2001. Structural characteristics of the patagium of *Ptychozoon kuhli* (Reptilia: Gekkonidae) in relation to parachuting locomotion. J. Morphol. 247, 252–263.

Siler, C.D., Oaks, J.R., Cobb, K., Ota, H., Brown, R.M., 2014. Critically endangered island endemic or peripheral population of a widespread species? Conservation genetics of Kikuchi’s gecko and the global challenge of protecting peripheral oceanic island endemic vertebrates. Divers. Distrib. 20, 756–772.

Siler, C.D., Oaks, J. R., Welton, L.J., Linkem, C.W., Swab, J.C., Diesmos, A.C., Brown, R.M., 2012. Did geckos ride the Palawan raft to the Philippines? J. Biogeogr. 39, 1217–1234.

Safonova, Y., Bankevich, A., Pevzner, P.A., 2015. dipSPAdes: assembler for highly polymorphic diploid genomes. J. Comp. Biol. 22, 528–545.

Smith, M.A., 1935. The Fauna of British India, Including Ceylon and Burma. Reptilia and Amphibia. II. Sauria. Taylor & Francis Ltd., London.

Snir, S., Rao, S., 2012. Quartet MaxCut: a fast algorithm for amalgamating quartet trees. Mol. Phylogenet. Evol. 62, 1–8.

Stejneger, L., 1902. *Ptychozoon kuhli*, a new name for *P. homalocephalus*. Proc. Biol. Soc. Washington 15, 37.

Stejneger, L. 1936. Types of the amphibian and reptilian genera proposed by Laurenti in 1768. Copeia 1936, 133–141.

Swofford, D.L., 2003. PAUP*: Phylogenetic Analysis Using Parsimony (and other methods). Sinauer Associates, Sunderland, Massachusetts, USA.

Tho, Y.P., 1974. Camouflage in the flying gecko, *Ptychozoon kuhli* Stejn. Malay. Nat. J. 28, 36.

Uetz, P., Freed, P. Jirí Hošek (eds.), The Reptile Database, http://www.reptile-database.org, accessed [November 2018].

Wermuth, H., 1965. Liste der rezenten Amphibien und Reptilien. Gekkonidae, Pygopodidae, Xantusiidae. Das Tierreich, 80, Walterv de Gruyter, Berlin, XXII + 246 pp.

Wilcox, T.P., Zwickl, D.J., Heath, T.A. Hillis, D.M., 2002. Phylogenetic relationships of the dwarf boas and a comparison of Bayesian and bootstrap measures of phylogenetic support. Mol. Phylogenet. Evol. 25, 361–371.

Vetter, R.S., Brodie, E.D. 1977. Background color selection and antipredator behavior of the flying gecko, *Ptychozoon kuhli*. Herpetologica 33, 464–467.

Wallach, V., Wuester, W., Broadley, D.G., 2009. In praise of subgenera: taxonomic status of cobras of the genus *Naja* Laurenti (Serpentes: Elapidae). Zootaxa 2236, 26–36.

Young, B.A., Lee, C.E., Daley, K.M., 2002. On a flap and a foot: aerial locomotion in the “Flying” Gecko, *Ptychozoon kuhli*. J. Herpetol. 36, 412–418.

Yuan, Z.Y., Zhou, W.W., Chen, X., Poyarkov Jr, N.A., Chen, H.M., Jang-Liaw, N.H., Chou, W.H., Matzke, N.J., Iizuka, K., Min, M.S., Kuzmin, S.L., 2016. Spatiotemporal diversification of the true frogs (genus Rana): a historical framework for a widely studied group of model organisms. Syst. Biol. 65(5), 824–842.

Zhang, C., Sayyari, E. Mirarab, S., 2017. Astral-III: increased scalability and impacts of contracting low support branches. In RECOMB International Workshop on Comparative Genomics, 53–75 (Springer, 2017).

